# Homeostatic regulation of intrinsic neuronal excitability in visual thalamic relay cells induced by brief monocular deprivation

**DOI:** 10.64898/2026.05.19.726212

**Authors:** Aurore Aziz, Laure Fronzaroli-Molinieres, Cécile Iborra, Maël Duménieu, Emilie Zanin, Thierry David, Danièle Denis, Juan José Garrido, Romain Brette, Michaël Russier, Dominique Debanne

## Abstract

Homeostatic plasticity of intrinsic excitability (IE) in the visual system has been essentially shown at the cortical level but whether thalamic nuclei also express homeostatic plasticity of IE is unknown. We show here that 4 days of monocular deprivation (MD) at eye opening induces a homeostatic change in IE in dorsal lateral geniculate nucleus (dLGN) neurons. Neurons recorded in the dLGN region activated by the deprived eye are more excitable than neurons recorded in the dLGN region activated by the open eye. No significant changes were observed following 7 days of MD, however. Enhanced excitability in neurons from the deprived side after 4 days of MD was associated with a reduced Kv1-dependent LTP-IE, a smaller voltage ramp, and a reduced inter-spike interval, suggesting that Kv1 channels are down-regulated in deprived dLGN neurons. Furthermore, the ankyrin G signal of the axon initial segment was larger in deprived dLGN neurons compared with open ones, indicating that Nav1 channel number also undergoes homeostatic regulation, and Kv1.1 channel signals were lower in deprived neurons compared to open ones. In addition, electrical coupling was found to be strengthened in neurons displaying enhanced IE following either brief (4 days) or long (10 days) MD. These results suggest that homeostatic and Hebbian plasticity in the dLGN share common expression mechanisms involving the regulation of Kv1 channels, Nav1 channels and electrical coupling between relay neurons.

## Introduction

Amblyopia is an acquired visual deficit resulting from abnormal visual experience during childhood and caused by strabismus, congenital cataract, anisometropia or ptosis which cannot be compensated by optic means at the adult stage (Holmes & Clarke, 2006). Amblyopia is generally attributed to the critical period of ocular dominance plasticity occurring during postnatal development that closes around sexual maturity in most species (Morishita & Hensch, 2008). First, observed in the visual cortex (Wiesel & Hubel, 1963; Hubel & Wiesel, 1970; Kleinschmidt *et al*., 1987; Frégnac *et al*., 1988; Nataraj *et al*., 2010), functional Hebbian-like changes in ocular dominance underlying amblyopia have more recently been also identified in the dorsal lateral geniculate nucleus (dLGN) (Jaepel *et al*., 2017; Sommeijer *et al*., 2017; Rose & Bonhoeffer, 2018). In particular, monocular deprivation for 10 days after eye opening has been shown to induce a reduction of IE in dLGN relay neurons found in zone activated by the deprived eye (Duménieu *et al*., 2025), indicating that visual stimulation might enhance IE. In addition, retinal input stimulation at 40 Hz during 10 minutes induces long term potentiation of intrinsic excitability (LTP-IE) in dLGN neurons (Duménieu *et al*., 2025). LTP-IE in thalamic relay neurons requires spiking activity, activation of L-type calcium channels for its induction and is mediated by the down-regulation of Kv1 channels.

Hebbian and homeostatic plasticity work hand-in-hand (Turrigiano, 2017). Homeostatic regulation of IE occurs to compensate chronic increases or decreases of network activity (Turrigiano & Nelson, 2004) and has been reported in neocortical neurons and hippocampal neurons *in vitro* (Desai *et al*., 1999; Karmarkar & Buonomano, 2006; Grubb & Burrone, 2010; Cudmore *et al*., 2010; Gasselin *et al*., 2015; Zbili *et al*., 2021). Homeostatic plasticity has been also reported *in vivo*. Visual cortical responsiveness is upregulated following brief monocular deprivation (MD) in both animal models (Maffei *et al*., 2004; Mrsic-Flogel *et al*., 2007) and humans (Lunghi *et al*., 2011, 2015; Binda *et al*., 2018), indicating that homeostatic plasticity is present in the visual cortex. However, the existence of homeostatic plasticity in the visual thalamus is much less documented (Van Hook *et al*., 2020; Bhandari *et al*., 2022). While homeostatic regulation of synaptic transmission has been reported in thalamic neurons (Krahe & Guido, 2011), homeostatic regulation of intrinsic plasticity has not been explored in dLGN neurons following MD.

We show here that MD for 4 days from eye opening enhances intrinsic neuronal excitability in neurons activated by the deprived eye. LTP-IE was reduced in neurons that are more excitable after 4 days of MD suggesting that enhanced IE is mediated by the reduction in Kv1.1 channel activity. In addition, neurons displaying an elevated IE after 4 days of MD display a reduced subthreshold voltage ramp and inter-spike interval. Ankyrin G staining was found to be larger and Kv1.1 labelling was found to be weaker in deprived compared to open neurons, confirming the electrophysiological characterization of homeostatic regulation. After 7 days of MD, no significant change in excitability is observed between neurons activated by the deprived or the open eye, but LTP-IE is significantly reduced on both sides. In addition, electrical coupling was found to be higher in neurons expressing enhanced IE following brief (4 days) and long (10 days) MD. These results indicate that homeostatic and Hebbian plasticity in the dLGN share common expression mechanisms involving the regulation of Kv1 channels and electrical coupling.

## Results

### Brief monocular deprivation homeostatically enhances IE in dLGN neurons

In order to determine the effects of brief monocular deprivation (MD) on neuronal excitability at the thalamic stage, Long-Evans rats were monocularly deprived just before eye opening (i.e., at P12) for 4 or 7 days and acute slices containing the dLGN were obtained (**Figure 1A**). Relay cells were recorded in whole-cell configuration from the monocular segment of the contralateral projection zone to the closed eye or from the monocular segment of the contralateral projection zone to the open eye of the dLGN. Compared to neurons activated by the open eye, neurons activated by the deprived eye were found to be more excitable after 4 days of monocular deprivation (MD) and exhibited a lower rheobase when the neuron was held at a membrane potential of −56 mV to inactivate the T-type calcium current (**Figure 1B** and **1C**). No significant difference in input resistance, nor in AP width, AP threshold and AP amplitude were observed between neurons from the open and deprived sides (**Figure S1A** & **S1B**). In contrast to what is observed after 4 days of MD, no significant change in excitability was observed between the open and deprived sides following 7 days of MD (**Figure 1D** & **1E**). The rheobase was significantly modified between the open and deprived sides following 4 days but not 7 days of MD (**Figure 1F**). Interestingly, when one considers on the same graph the data obtained after 10 days of MD from a previously published paper (Duménieu *et al*., 2025), the rheobase evolves from homeostatic change after 4 days of MD to Hebbian-like change following 10 days of MD (**Figure 1F**). The excitability level and rheobase of dLGN neurons recorded in control animals of the same age (i.e., with binocular vision), were found to be intermediate between those of deprived side and those of open side (**Figure S2**).

**Figure 1.**
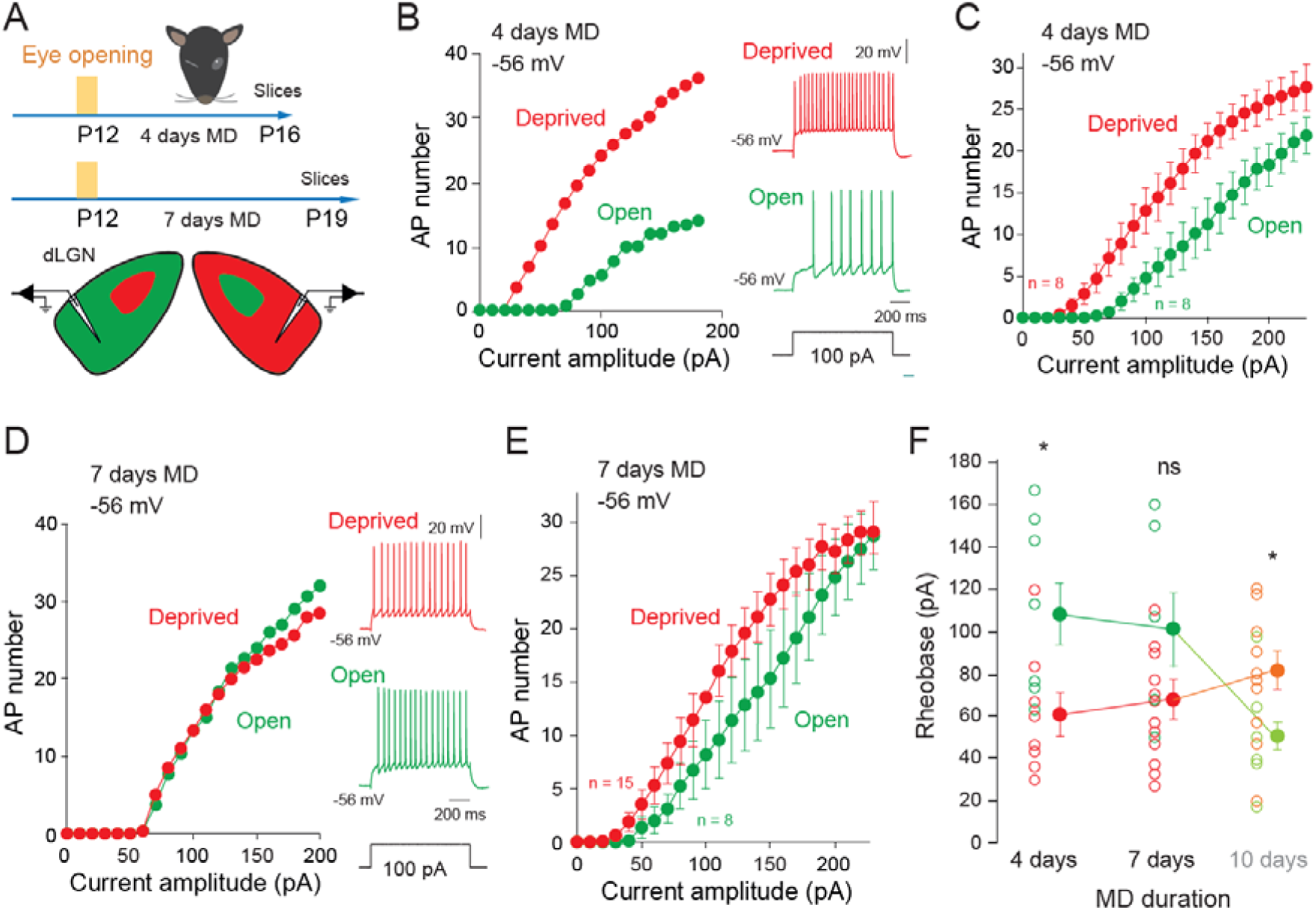
dLGN neurons express homeostatic plasticity of IE following 4 days of MD. A. Experimental protocols. B. Representative input-output curves of neurons recorded in the deprived zone (red) and in the open zone (green) at −56 mV following 4 days of MD. C. Group data showing a clear separation between deprived and open neurons. D. Representative input-output curves of neurons recorded in the deprived zone (red) and in the open zone (green) at −56 mV following 7 days of MD. E. Group data showing overlapping input-output curves of deprived and open neurons. F. Rheobase changes as a function of MD duration. Note the homeostatic regulation at 4 days of MD and Hebbian-like regulation at 10 days of MD. The data at 10 days comes from Duménieu et al., 2025. *, Mann-Whitney U-test, p<0.05. ns, p>0.1.

The same difference between the open and deprived sides was observed at −65 mV following 4 days of MD (**Figure S3A**), thus confirming the homeostatic change in IE observed at −56 mV. Nevertheless, a significant difference in the rheobase was only seen when the rheobase was measured without the contribution of T-type calcium potential by selecting in the analysis the inflexion point (**Figure S3B**) or by using the selective antagonist of T-type calcium current, TTAP2 **(Figure S3C**). No significant difference between the open and deprived sides was also observed at −65 mV following 7 days of MD (**Figure S4**), thus confirming the observation made at −56 mV.

### Reduced amplitude of LTP-IE after 4 and 7 days of MD

To determine the relationship between homeostatic MD-induced changes in IE with Hebbian-like plasticity of IE already reported (Duménieu *et al*., 2025), we checked whether the amplitude of LTP-IE induced by trains of spiking activity is altered after MD. First, we determined the magnitude of LTP-IE in rats that had normal visual experience at P16 and P19 (i.e., at ages corresponding to 4 and 7 days of MD, respectively). LTP-IE was induced by spiking activity at 10 Hz during 10 minutes (**Figure S5A**). As previously reported (Duménieu *et al*., 2025), the amplitude of LTP-IE increased from P16 to P19 (148 ± 10%, n = 7 at P16 and 240 ± 22%, n = 15 at P19; **Figure S5B** and **S5C**). After 4 days of MD, the magnitude of LTP-IE was found to be significantly reduced in neurons activated by the deprived eye (i.e., neurons that expressed an enhanced neuronal excitability; **Figure 2A** and **2C**) while neurons activated by the open eye expressed the same level of LTP-IE (**Figure 2A** and **2C**). After 7 days of MD, the magnitude of LTP-IE was significantly reduced in neurons activated by both the open and deprived eye (**Figure 2B** and **2D**). After 10 days of MD, the magnitude of LTP-IE has been shown to be reduced (**Figure S6**) in neurons activated by the open eye (Duménieu *et al*., 2025). Thus, LTP-IE is reduced on the deprived eye until 10 days of MD whereas on the open eye, LTP-IE is reduced from 7 days of MD (**Figure 2E**). These data suggest that homeostatic and Hebbian-like plasticity in dLGN neurons are expressed through the modulation of Kv1 channel activity.

**Figure 2.**
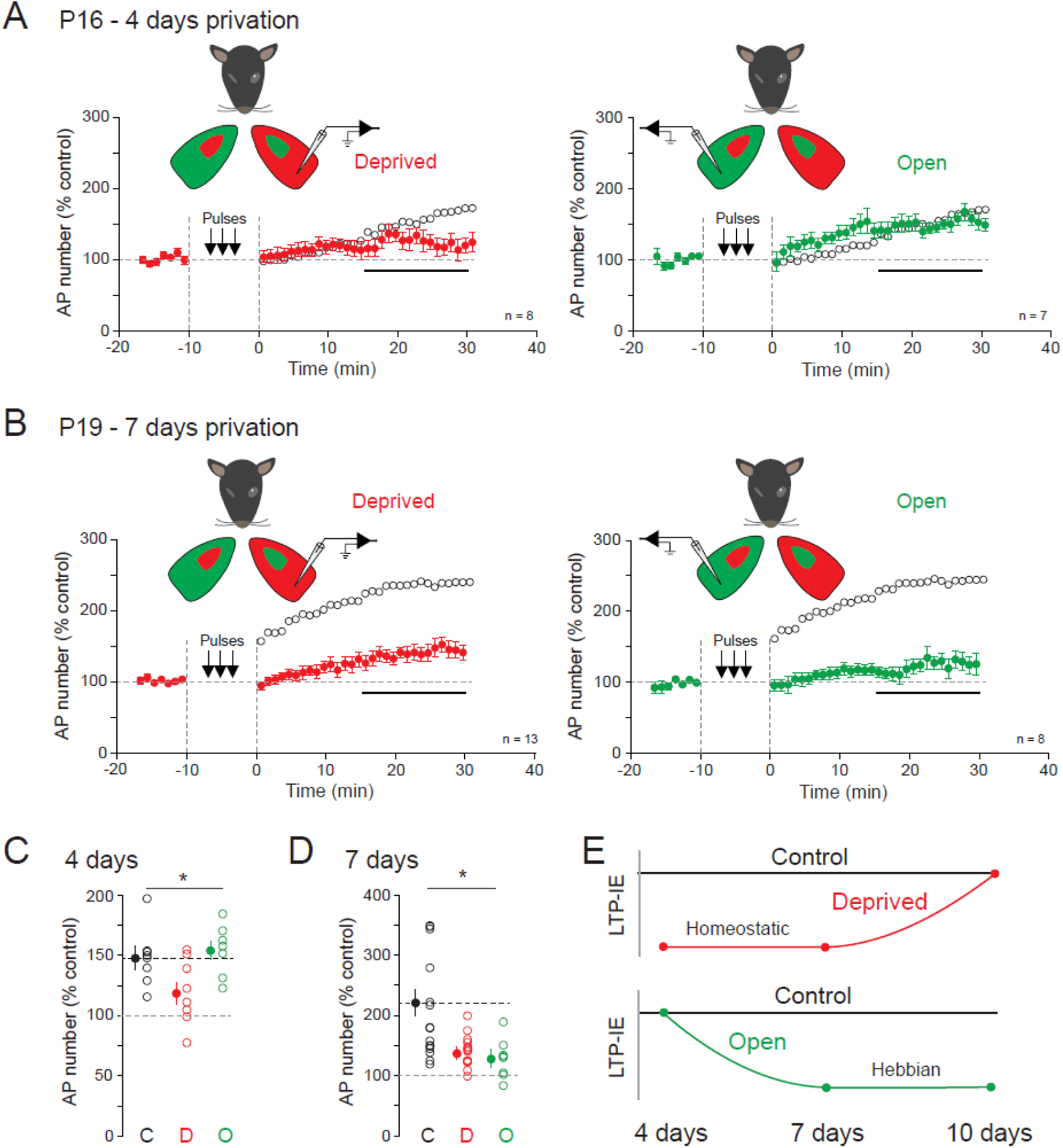
Modulation of LTP-IE after 4 and 7 days of MD. A. LTP-IE induced in neurons activated by the deprived eye (left) and the open eye (right) after 4 days of MD. Note the reduced magnitude in neurons activated by the deprived compared to the control (open black circles). B. LTP-IE induced in neurons activated by the deprived eye (left) and the open eye (right) after 7 days of MD. Note the reduction on both sides. C. Comparison of LTP-IE in control (C, P16), deprived (D, 4 days) and open (O, 4 days) neurons. *, Kruskal-Wallis test, p<0.05. D. Comparison of LTP-IE in control (C, P19), deprived (D, 7 days) and open (O, 7 days) neurons. *, Kruskal-Wallis test, p<0.05. E. Summary of the changes in LTP-IE after 4, 7 and 10 days. Homeostatic reduction of LTP-IE occurs on the deprived side after 4 and 7 days of MD while Hebbian reduction of LTP-IE occurs on the open side after 7 and 10 days of MD.

### Change in ISI and voltage ramp following 4 days of MD

LTP-IE expression is mediated by the downregulation of Kv1 channels (Duménieu *et al*., 2025). The reduction in the LTP-IE magnitude in neurons activated by the deprived eye and expressing an elevation in IE suggests that Kv1 channels are downregulated in these neurons. We therefore analyzed the electrophysiological cues of a differential change in Kv1 channel activity between deprived and open neurons after 4 days of MD. Kv1 channel inactivation is responsible for a subthreshold depolarizing voltage ramp that increases the inter-spike interval (ISI) of the first 2 APs compared to that of the second and third AP (Cudmore *et al*., 2010; Duménieu *et al*., 2025). We therefore analyzed these two parameters in dLGN neurons following 4 days of MD. The ISIs were found to be differentially modified in neurons activated by the deprived and open eyes (**Figure 3A**). A significant reduction of ISI2/ISI1 was found in open compared to deprived neurons (**Figure 3B**). In parallel, a clear ramp was visible on neurons activated by the open eye but not by the deprived eye (**Figure 3A** and **3C**). These data support the reduction of Kv1 in deprived vs. open neurons following 4 days of MD, further suggesting that similar mechanisms underlie both homeostatic and Hebbian plasticity.

**Figure 3.**
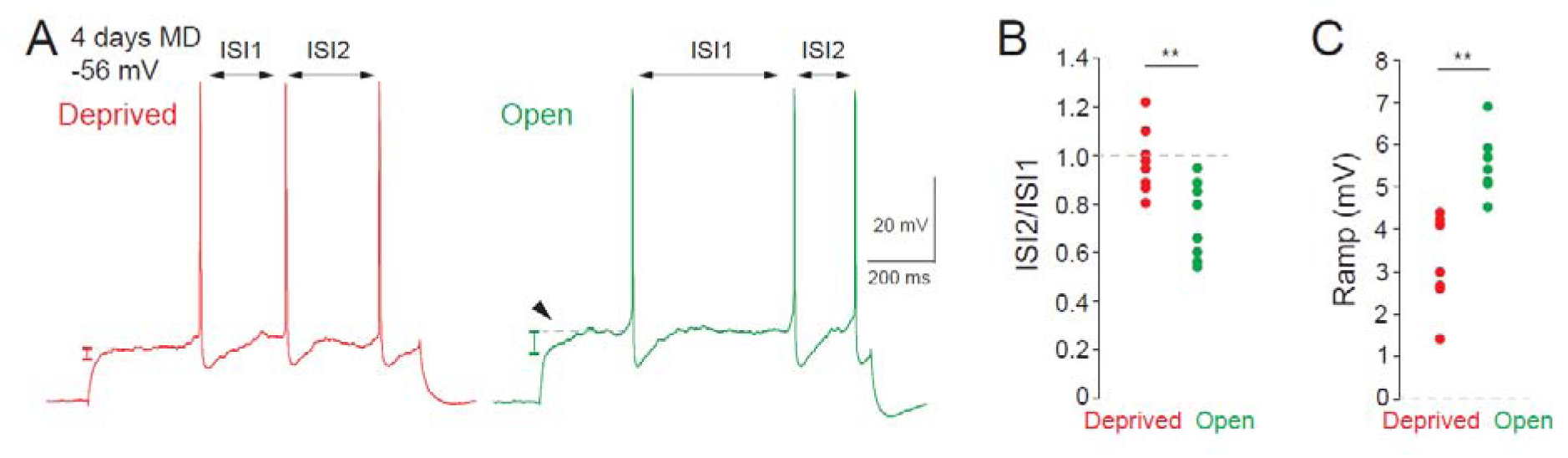
Modified inter-spike interval (ISI) and ramp amplitude after 4 days of MD. A. The first two ISIs are equal in neurons activated by the deprived eye but not in neurons activated by the open eye where ISI1 is much longer than ISI2. A voltage ramp immediately after the depolarizing onset is visible in neurons activated by the open but not the deprived eye. B. Group data of the ratio of ISI2/ISI1 for deprived and open dLGN neurons. **, Mann-Whitney U-test, p<0.01. C. Group data of the ramp amplitude for deprived and open dLGN neurons. **, Mann-Whitney U-test, p<0.01.

### Enhanced ankyrin G labeling and reduced Kv1.1 channel staining following MD

Next, we made double immunostaining of ankyrin G and Kv1.1 to see whether these proteins are differentially regulated following MD for 4 days. Ankyrin G labeling peaked at ∼9-10 µm and was found to be significantly larger on the deprived side following 4 days of MD compared to the open side, with a significant difference in the distal part of the ankyrin G peak (8-28 µm) (**Figure 4A**), suggesting that Nav channels density is larger in deprived versus open neurons. However, the AIS length was not significantly different, although the mean length was higher in deprived neurons (deprived: 30.2 ± 0.4 µm, n = 135; open: 28.8 ± 0.4 µm, n = 136; **Figure 4B**). The peak of Kv1.1 immunoreactivity was found to be shifted toward the end of the AIS (∼20 µm) as already reported (Kuba *et al*., 2015). Importantly, the Kv1.1 immunosignal was significantly weaker in deprived compared to open neurons (mean Kv1.1 signal: 70.7 ± 1.2 n = 127 in deprived neurons vs. 76.9 ± 1.6, n = 126 in open neurons; **Figure 4C**), in particular in the proximal region of the AIS (0-20 µm), thus confirming the electrophysiological data.

**Figure 4.**
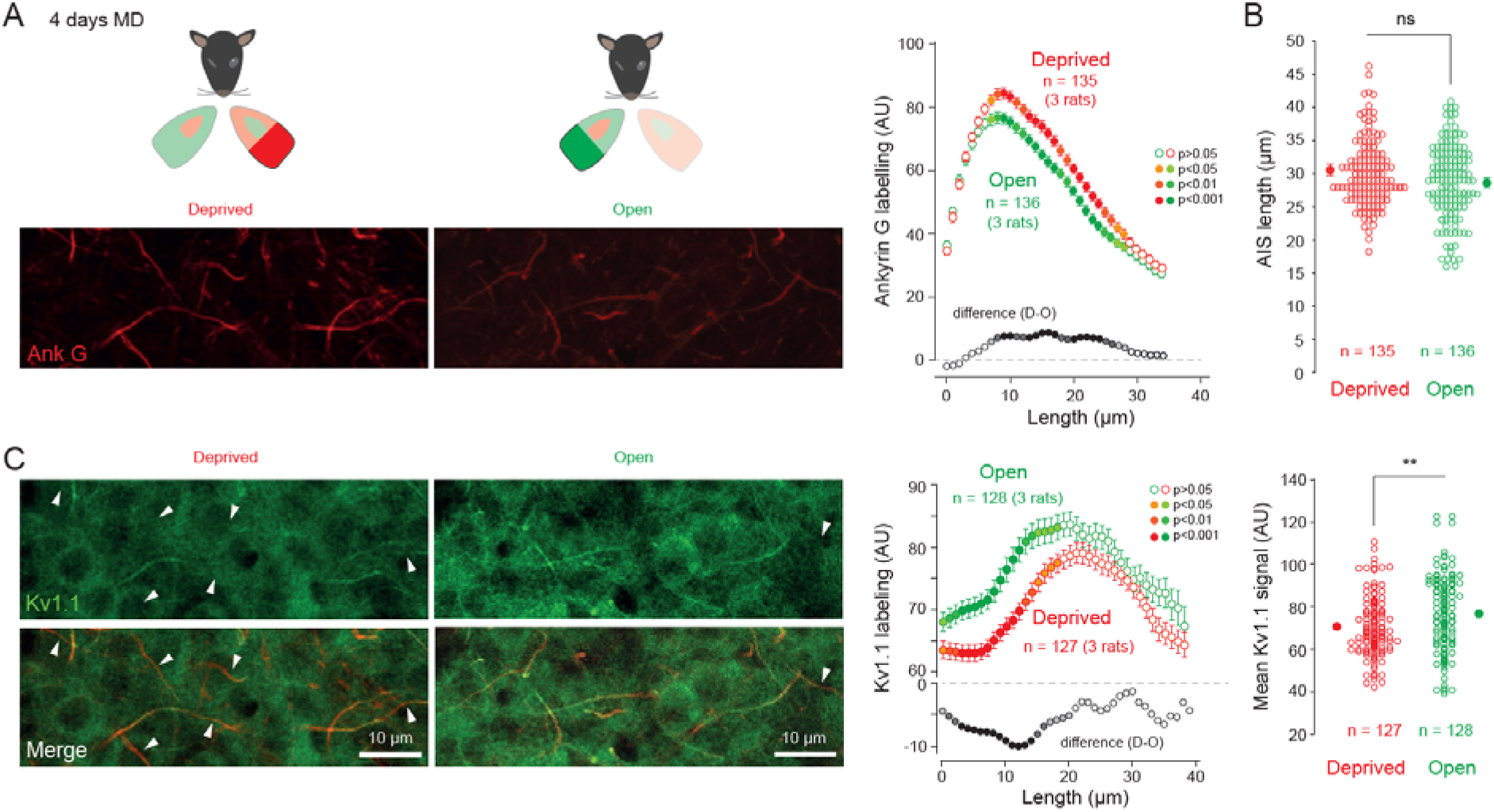
Opposite changes in ankyrin G and Kv1.1 staining following 4 days of MD. A. Left, immunostaining of ankyrin G from deprived neurons (left panel) and open ones (right panel). Right, comparison of the fluorescence signals from the AIS of deprived (red) and open (green) dLGN neurons. Note the significant difference near the pic of signal (8-27 µm; light color, Mann-Whitney U-test, p<0.05; medium color, Mann-Whitney U-test, p<0.01; plain color, Mann-Whitney U-test, p<0.001). B. Comparison of the length of AIS. ns, Mann-Whitney U-test, p>0.1. C. Left, immunostaining of Kv1.1 from deprived neurons (left panel) and open ones (right panel). White arrows indicate AIS with faint Kv1.1 labelling. Note the high number of white arrows in deprived vs. open side. Middle, comparison of the Kv1.1 staining in AIS from deprived and open neurons. Note the significant difference in the proximal region of the AIS (0-20 µm; light color, Mann-Whitney U-test, p<0.05; medium color, Mann-Whitney U-test, p<0.01; plain color, Mann-Whitney U-test, p<0.001). Right, comparison of the total signal Kv1.1 for each AIS. **, Mann-Whitney U-test, p<0.01.

### Change in ISI and voltage ramp following 7 days of MD

The reduction of LTP-IE on both deprived and open sides after 7 days of MD suggests that Kv1 channel activity is reduced in both cases compared to the control condition. We therefore examined whether the ISI and voltage-ramp are changed in control and MD dLGN neurons. Compared with controlled neurons, both deprived and open neurons after 7 days of MD displayed a higher ISI2/ISI1 ratio (**Figure 5A** and **5B**). In addition, the amplitude of the voltage ramp just before the first AP was significantly reduced (**Figure 5A** and **5C**). Both measures strongly argue for a reduction in Kv1 channel activity in both deprived and open neurons after 7 days of MD.

**Figure 5.**
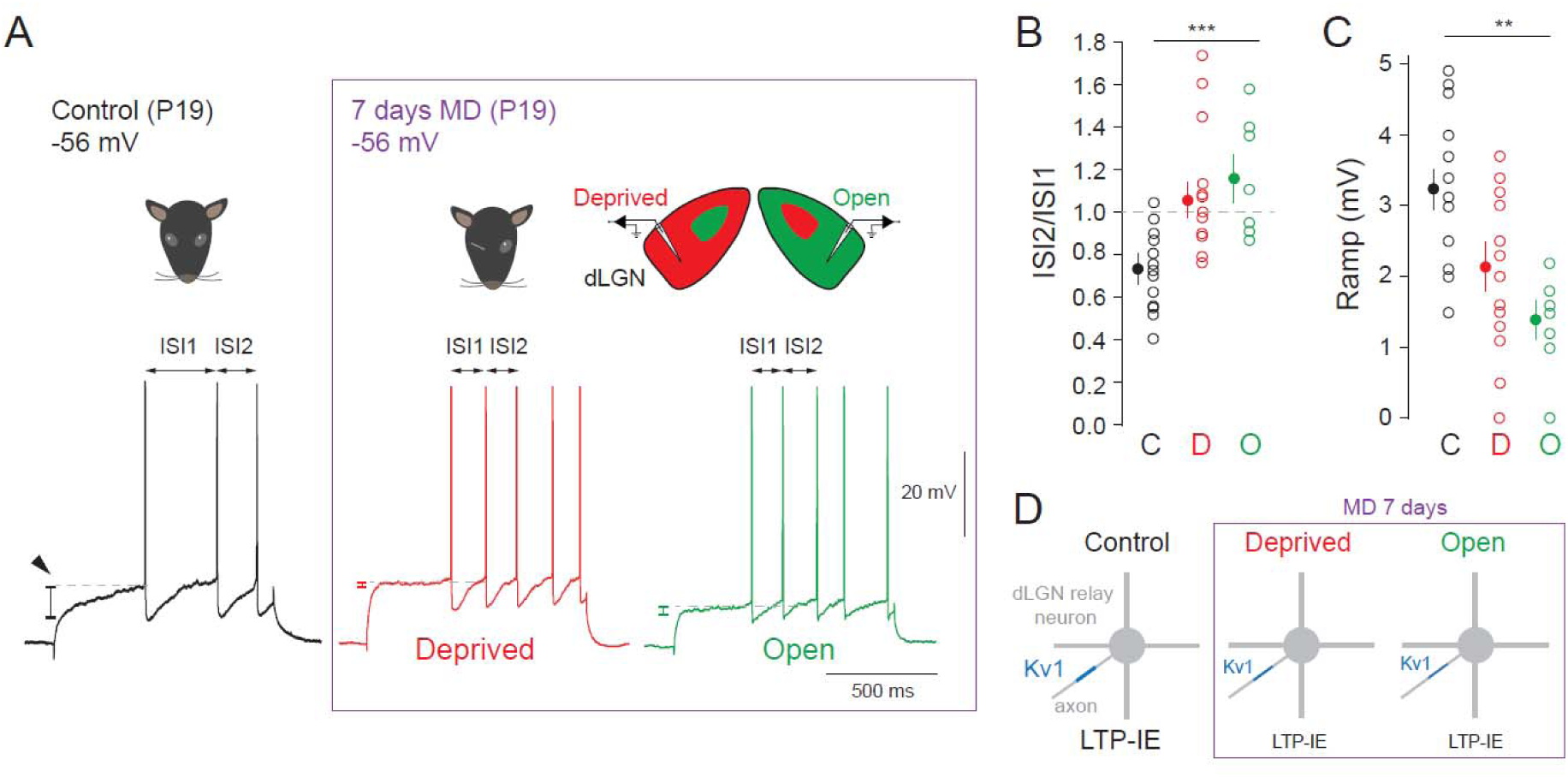
MD for 7 days augments the ISI and reduces the depolarizing voltage ramp. A. Representative voltage profile of a control neuron (i.e., from an animal with normal visual experience) in black, and voltage profiles of a neuron from the deprived dLGN region (red) and from the open region following 7 days of MD. B. Analysis of the ISI2/ISI1 ratio in control (black), deprived (green) and open (green) dLGN neurons. ***, Kruskal-Wallis test, p<0.001. C. Analysis of the voltage ramp in control (black), deprived (green) and open (green) dLGN neurons. **, Kruskal-Wallis test, p<0.01. D. Summary of the changes occurring following 7 days of MD. Kv1 channel density could be downregulated in both deprived and open dLGN neurons, thus accounting for the reduced LTP-IE.

### Modification in electrical coupling following MD

In the presence of synaptic blockers, transient hyperpolarizing potentials of constant amplitude were evoked by subthreshold depolarizations (**Figure 6A**). Their latency was found to be inversely proportional to the current injection (**Figure 6B**). In some cases, a small spikelet followed by a prominent hyperpolarization corresponding to the after-hyperpolarizing potential (AHP) of the AP was observed (**Figure S7A**) as previously shown in some electrical coupling (Galarreta & Hestrin, 2001; Dugué *et al*., 2009; Alcamí & Pereda, 2019). The spikelet derivative is roughly homothetic of the full AP derivative recorded in the same cell (**Figure S7B**). Electrical coupling disappeared in the presence of carbenoxolone (100 µM; **Figure S7C**) and declined during development (**Figure S7D**). Importantly, the electrical coupling was found to change upon MD. After 4 days of MD, coupled AHP were larger in deprived neurons (i.e., in excitable neurons) compared to open ones while after 10 days of MD the opposite was observed (**Figure 6C**). This reduced coupling after 10 days of MD in deprived neurons was further reinforced by the smaller proportion (∼30 vs. ∼50%) of coupling (**Figures S7E**). Altogether, homeostatic and Hebbian increases in IE are both associated with an enhanced electrical coupling.

**Figure 6.**
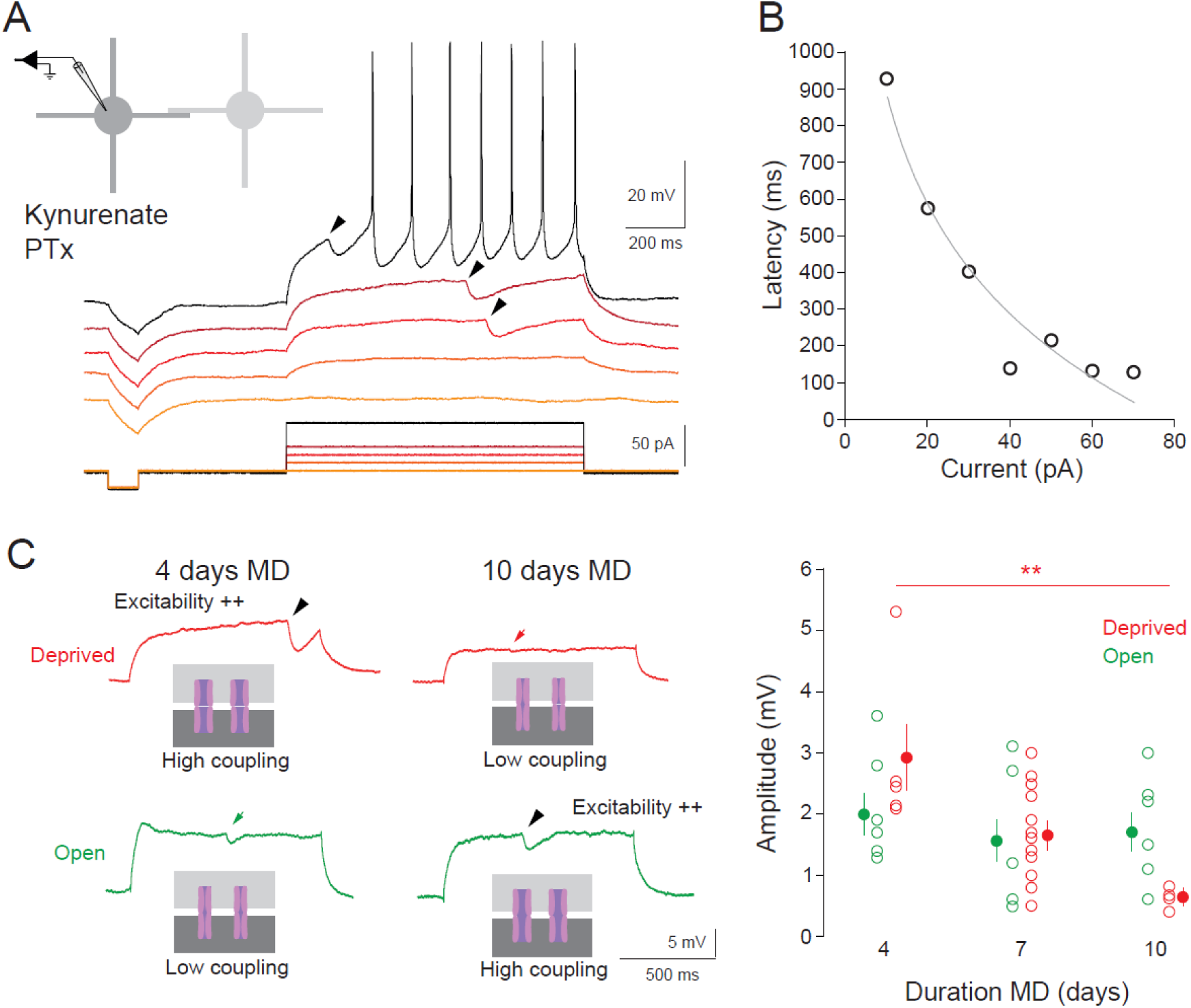
Changes in electrical coupling following MD. A. AHP transmitted by electrical coupling. In the presence of the synaptic blockers kynurenate and PTx, subthreshold hyperpolarizing events of constant amplitude are observed (arrow heads). B. Plot of the latency of hyperpolarizing events as a function of the current amplitude. Line corresponds to the fitted curve of the data (y = 427.6Ln(x) + 1867.2; R^2^=0.94). C. Coupling amplitude is modified following MD. At 4 days of MD (left column), the event amplitude is larger (high coupling) in neurons activated by the deprived eye compared to the open eye (low coupling). At 10 days of MD (right column), the event amplitude is larger (high coupling) in neurons corresponding to the open eye compared to the deprived eye (low coupling). Right graph, event amplitude as a function of MD duration. **, Kruskal-Wallis test, p<0.01.

## Discussion

We report here the existence of homeostatic plasticity of IE in dLGN relay neurons following brief MD of 4 days but not 7 days. After 10 days of MD, IE follows a Hebbian-like plasticity. Kv1-dependent LTP-IE is reduced in dLGN neurons expressing an enhanced IE following 4 days of MD, suggesting that homeostatic enhancement of IE is mediated by Kv1 channels and that homeostatic plasticity shares common expression mechanisms with Hebbian-like plasticity. Importantly, the enhanced IE after 4 days of MD is also associated with a smaller depolarizing voltage ramp, and a shorter first ISI, confirming the down-regulation Kv1 (**Figure 7**). Ankyrin G staining is larger while Kv1.1 labelling is weaker in deprived neurons compared to open neurons after MD for 4 days. LTP-IE is also reduced on both sides following 7 days of MD. Electrical coupling in dLGN neurons were found to be modified by MD with enhanced coupling in neurons displaying an elevated IE after 4 or 10 days of MD (**Figure 7**). Taken together, our data suggest that homeostatic and Hebbian plasticity in the dLGN share common expression mechanisms involving the regulation of Kv1 channels and electrical coupling.

**Figure 7.**
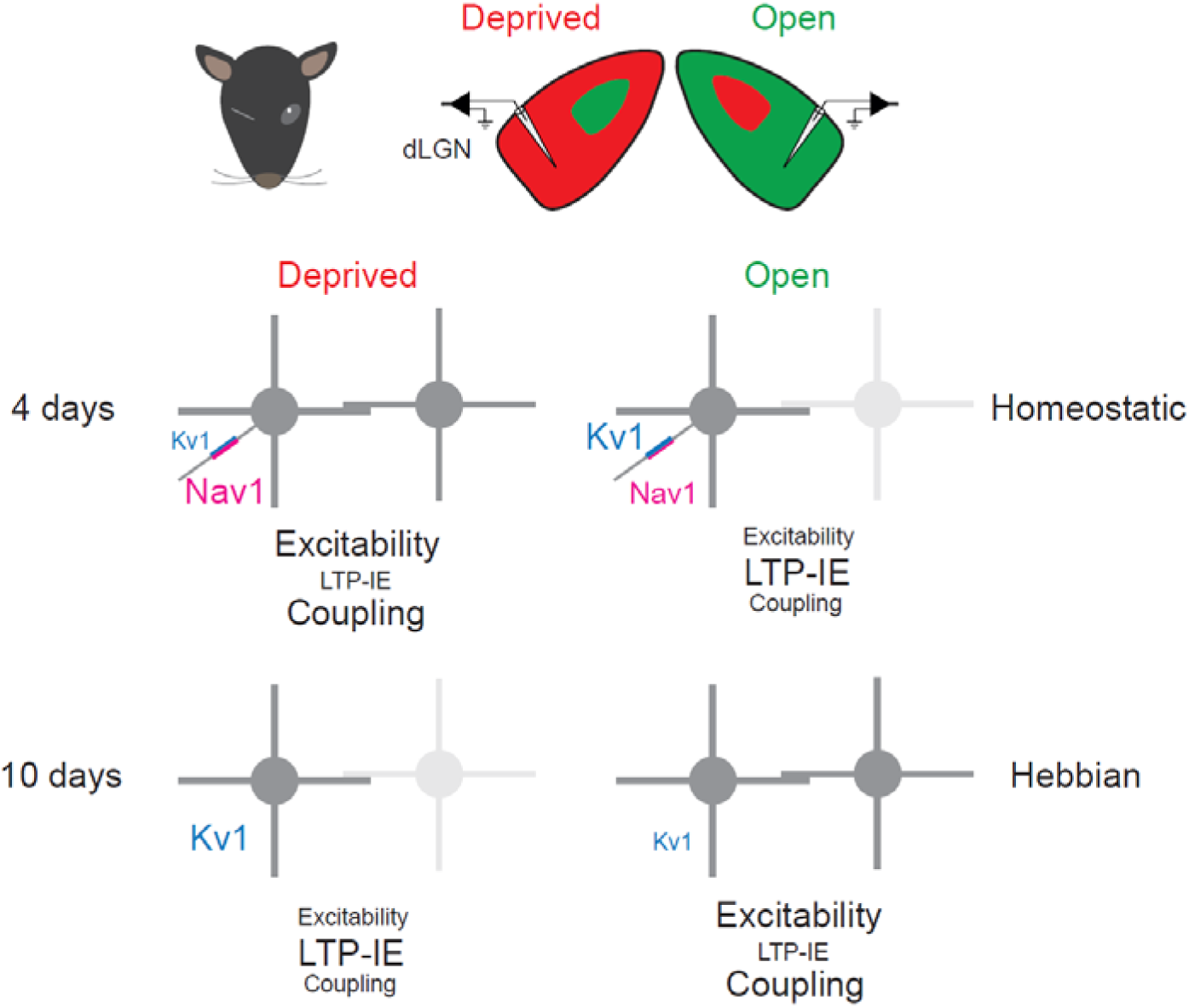
Summary of changes occurring in dLGN relay neurons following MD. At 4 days of MD, deprived neurons undergo a reduction in Kv1 channel density at the AIS accounting for both the increased excitability and the reduced Kv1-dependent LTP-IE. Parallelly, the density of Nav1 channels is slightly enhanced in deprived compared to open neurons. In open neurons, Kv1 channel level remains high allowing induction of LTP-IE. Electrical coupling is high in deprived neurons (most excitable) compared to the open ones. At 10 days, open dLGN neurons undergo a downregulation in Kv1 channel density accounting for both the increased excitability and the reduced Kv1-dependent LTP-IE. Electrical coupling is high in open neurons (most excitable) compared to the deprived ones.

### Homeostatic plasticity of IE in dLGN neurons

We show here that reducing visual activity for 4 days induces a differential change in IE in dLGN in favor of the deprived eye. This compensatory elevation in IE is not observed after 7 days of MD, however. Homeostatic plasticity of IE in dLGN neurons has been observed following experimental glaucoma (Van Hook *et al*., 2020) or following binocular enucleation ((Bhandari *et al*., 2022). However, in these reports, the underlying mechanisms of expression were not investigated. Our data indicate that regulation of Kv1 channel is a key parameter in homeostatic elevation of IE.

In parallel with Kv1 changes, the AIS length was found to be longer and the ankyrin-G signal stronger in deprived dLGN neurons compared to open dLGN neurons. Nav channels labeling generally match ankyrin-G labeling (Garrido *et al*., 2003; Jenkins & Bender, 2025). Thus, an augmentation of Nav1 channels is expected to occur in parallel with the reduction of Kv1 channels as already reported (Zbili *et al*., 2021).

Homeostatic plasticity in the visual cortex has been observed in humans following brief monocular deprivation (Lunghi *et al*., 2011, 2015; Binda *et al*., 2018). While the mechanism may involve intracortical plasticity, this phenomenon could also result from enhanced neuronal excitability of deprived LGN neurons. Homeostatic plasticity is classically thought to result from persistent changes in network activity whereas Hebbian plasticity is thought to result from brief changes in neural activity. This view has recently been modified with reports showing rapid homeostatic plasticity (Riegle & Meyer, 2007; Ibata *et al*., 2008; Morgan *et al*., 2019; Jamann *et al*., 2021). We show here that homeostatic plasticity also precedes Hebbian plasticity in dLGN neurons. Such temporal order has already been observed in the visual cortex following MD (Mrsic-Flogel *et al*., 2007)

### Common mechanisms for Hebbian and homeostatic plasticity

We previously showed that LTP-IE induced by retinal inputs at 40 Hz is expressed through the down-regulation of Kv1 channels (Duménieu *et al*., 2025). In fact, following induction of LTP-IE, the AP threshold was hyperpolarized, the latency of the first regular AP was reduced and above all, LTP-IE was absent in the presence of the specific Kv1 channel blocker, dendrotoxin. Following 10 days of MD, LTP-IE was reduced in neurons activated by the open eye that expressed a Hebbian-like elevated IE. We show here that LTP-IE is also reduced in neurons from the deprived eye that expressed a homeostatic elevation in IE following 4 days of MD. Thus, LTP-IE is modified equally in Hebbian-like or homeostatic regulation of IE.

The contribution of Kv1 channels has already been reported in the context of homeostatic plasticity. Activity deprivation induced by treatment for 48-72 h with antagonists of glutamate receptor reduces the Kv1-mediated current in hippocampal CA1 pyramidal neurons (Cudmore *et al*., 2010; Zbili *et al*., 2021). Kv1 channels are replaced by Kv7 channels following auditory deprivation (Kuba *et al*., 2015).

### Regulation of both ankyrin G and Kv1.1 channel density

We show here that following 4 days of MD, the ankyrin G immunolabeling is significantly enhanced in the AIS of deprived neurons compared to open neurons while the Kv1.1 staining was lower in the AIS of deprived neurons compared to open ones. The increase in ankyrin G in deprived neurons suggests that Nav1 channels are significantly upregulated in these neurons. The reduction of Kv1.1 labeling in deprived neurons confirms the electrophysiological findings: 1) reduced Kv1-dependent LTP-IE on deprived side, 2) reduced depolarizing ramp and 3) equilibrated ISI. The coregulation of at least two ion channels has already been reported following *in vitro* and *in vivo* manipulations of neuronal activity (Desai *et al*., 1999; Kuba *et al*., 2015) and could be a general principal in regulation of intrinsic neuronal excitability (Marder & Goaillard, 2006).

### Reduced LTP-IE following 7 days of MD

We report here a reduced LTP-IE following 7 days of MD initiated at eye opening. While, no significant changes in IE have been observed, the magnitude of LTP-IE is reduced on both the deprived and open side. The reduction of LTP-IE on the deprived side could be due to the homeostatic reduction of Kv1 channel activity while the reduction of LTP-IE on the open side could correspond to the Hebbian-reduction of Kv1 channel activity. This assumption is supported by the fact that the ISI ratio and the voltage ramp were significantly modified in deprived. In this view, however, the global excitability should be enhanced on both sides compared to the control. This is not the case, however. Therefore, the bilateral reduction in Kv1 channel activity should be compensated by the homeostatic and Hebbian shortening of the AIS.

If Kv1 channels are reduced following 7 days of MD on both sides, why their global excitability is comparable to control neurons? This could be possible if one assumes that the increase in excitability induced by the reduction of Kv1 channel activity is compensated by a reduction in Nav channels in MD neurons for 7 days.

### Regulation of electrical coupling following 4 days and 10 days of MD

We identified electrical coupling in rat dLGN relay neurons. Electrical coupling has been reported in ventrobasal nucleus (VBN) relay cells in developing mice (Lee *et al*., 2010). Gap junctions between relay neurons have already been reported in the cat LGN where they help oscillation synchrony (Hughes *et al*., 2004, 2011). Our data suggest that electrical coupling between dLGN neurons varies with visual experience. MD-induced enhancement of IE is associated with an larger electrical coupling both after 4 days or 10 days of MD. Long lasting regulation of electrical synapses have already been reported in thalamic reticular neurons following manipulation of neuronal activity (Landisman & Connors, 2005; Haas *et al*., 2011; Sevetson *et al*., 2017; Fricker *et al*., 2021; Vaughn *et al*., 2025). However, the regulation of gap junctions following MD has never been reported so far.

The electrical coupling between dLGN neurons mostly transmit AHPs. This observation is consistent with previous observations made in neocortical, hippocampal and cerebellar neurons (Galarreta & Hestrin, 2001; Zsiros & Maccaferri, 2005; Dugué *et al*., 2009; Alcamí & Pereda, 2019). At glance, electrical transmission of AHPs would reduce spiking excitability. But the transmitted AHP could also avoid inactivation of Nav channels during intense spiking frequency as observed in deprived neurons following 4 days of MD or in open neurons after 10 days of MD.

dLGN neurons are generally considered as oscillatory neurons as they display strong oscillatory activity in the theta frequency range during sleep (Krosigk von *et al*., 1993; Steriade *et al*., 1993) and in the gamma frequency range during awakening (Storchi *et al*., 2017). Increasing electrical coupling has been shown to modulate the frequency of oscillator usually towards lower frequency (Kepler *et al*., 1990).

## Methods

### Monocular deprivation

All experiments were conducted according to the European and Institutional guidelines (Council Directive 86/609/EEC and French National Research Council and approved by the local health authority (Veterinary services, Préfecture des Bouches-du-Rhône, Marseille; APAFIS n°7661-2016111714137296 v4)). Young (P12-13) Long Evans rats of both sexes were anesthetized with isoflurane and lids of the right eye were sutured for 4 to 7 days after adding eye drops containing a local anaesthetic (tetracaine 1%). To improve the persistence of sutures and therefore to reduce the number of animals, a portion of eye-patch adjusted to the size of the rat head was glued with surgical glue onto the sutured eye (**Figure S8**). Sutures were checked each day and if not intact, animals were not used.

### Ex-vivo slices of rat dLGN

Thalamic slices (350 µm) were obtained from 16- to 19-day-old Long Evans rats of both sexes. Rats were deeply anesthetized with isoflurane and killed by decapitation. Slices were cut in an ice-cold solution containing (in mM): 92 *n*-methyl-D-glutamine, 30 NaHCO_3_, 25 D-glucose, 10 MgCl_2_, 2.5 KCl, 0.5 CaCl_2_, 1.2 NaH_2_PO_4_, 20 Hepes, 5 sodium ascorbate, 2 thiourea, and 3 sodium pyruvate and bubbled with 95% O_2_ - 5% CO_2_ (pH 7.4). Slices recovered (20-30 min) in the NMDG solution before being transferred in a solution containing (in mM): 125 NaCl, 26 NaHCO_3_, 2 CaCl_2_, 2.5 KCl, 2 MgCl_2_, 0.8 NaH_2_PO_4_, and 10 D-glucose and equilibrated with 95% O_2_ - 5% CO_2_. Each slice was transferred to a submerged chamber mounted on an upright microscope (Olympus, BX51 WI) and neurons were visualized using differential interference contrast infrared video-microscopy.

### Electrophysiology

Whole-cell patch-clamp recordings were obtained from relay neurons in the contralateral projection zone of the dLGN. The external saline contained (in mM): 125 NaCl, 26 NaHCO_3_, 3 CaCl_2_, 2.5 KCl, 2 MgCl_2_, 0.8 NaH_2_PO_4_, and 10 D-glucose and equilibrated with 95% O_2_ - 5% CO_2_. Synaptic inhibition was blocked with 100 µM picrotoxin and excitatory synaptic transmission was blocked with 2 mM kynurenate. Patch pipettes (5-10 MΩ) were pulled from borosilicate glass and filled with an intracellular solution containing (in mM) 120 K-gluconate, 20 KCl, 10 Hepes, 0.5 EGTA, 2 MgCl_2_, 2 Na_2_ATP and 0.3 NaGTP (pH 7.4). Recordings were performed with a MultiClamp-700B (Molecular Devices) at 30°C in a temperature-controlled recording chamber (Luigs & Neumann, Ratingen, Germany). The membrane potential was not corrected for the liquid junction potential (−13 mV). In all recordings, access resistance was fully compensated using bridge balance and capacitance neutralization (>70%). dLGN neurons were recorded in current-clamp and input resistance was monitored throughout the duration of the experiments. Cells that display a variation >20% were discarded from final analysis. Voltage and current signals were low pass-filtered (10 kHz) and sequences of 2 s were acquired at 20 kHz with pClamp (Axon Instruments, Molecular Devices).

### Excitability measure after monocular deprivation

The excitability of dLGN neurons after monocular deprivation was tested at two different membrane potentials: −65 mV that approximately corresponds to the resting membrane potential of dLGN neurons and at −56 mV that corresponds to the transmission mode without bursting observed during wakefulness (Weyand *et al*., 2001). Steps of depolarizing current with an increment of 10 pA were injected and the number of AP was counted for each current value. The rheobase was determined as the smallest current that elicits at least an AP.

### LTP-IE induction

LTP-IE was induced as previously reported (Russier *et al*., 2024; Duménieu *et al*., 2025). IE was tested with injection of current pulses (150-250 nA, 800 ms) to elicit 3-4 action potentials in control conditions at a frequency of 0.1 Hz. The amplitude of the current pulse and the membrane potential (−65 mV) were kept constant before and after induction of LTP-IE. LTP-IE was quantified 15-30 minutes after the end of the induction protocol.

### Data analysis

Electrophysiological signals were analysed with ClampFit (Axon Instruments, Molecular Devices). Spikes were counted using Igor Pro software (Wavemetrics). Input resistance was calculated from voltage response to small negative current pulses (typically, −20 pA, 250 ms). LTP-IE was measured over a period of 10 minutes, 20 minutes after the beginning of the post-stimulation period.

Electrical coupling was identified in dLGN neurons as hyperpolarization of constant amplitude preceded in some cases by a spikelet. Data from Duménieu et al., have been analysed for the presence of electrical coupling.

### Drugs

All chemicals were bath applied. Picrotoxin was purchased from Abcam (UK), carbenoxolone from Tocris Bioscience (UK), kynurenate from Sigma-Aldrich (USA), and 3,5-dichloro-N-[1-(2,2-dimethyl-tetrahydro-pyran-4-ylmethyl)-4-fluoro-piperidin-4-ylmethyl]-benzamide (TTA-P2) from Alomone Labs (Israel).

### Immunostaining

Long Evans rats monocularly deprived for 4 days were deeply anesthetized with isoflurane until death and then perfused with ice-cold 4% paraformaldehyde diluted in glucose 20 mM in phosphate buffer 0.1 M pH 7.4. Brains were removed from the skull and post-fixed overnight at 4°C in the same fixative solution. 70 µm thick sections were cut using a Leica VT1200S vibratome and then processed for immuno-histochemical detection. Brain slices were washed in DPBS (Gibco #14190-094) and incubated 30 min in sodium citrate 10 mM pH 8.5 at 80°C for antigen retrieval. Then, the slices were washed with phosphate buffer 0.1 M pH 7.4 (PB). Immunodetection was done in free-floating sections. Brain slices were treated with 50 mM NH_4_Cl for 30 min and incubated in blocking buffers to diminish nonspecific binding, using in a first step a PB solution containing 1% (w/v) BSA (Sigma #A3059), 0.3% (v/v) TritonX100 (Merk #648463) for 30 min, then a PB solution containing 5% (w/v) BSA, 0.3% (v/v) Triton X-100 for 2 h. Slices were then incubated overnight at 4°C with primary antibodies, mouse anti Kv1.1 clone K36/15 IgG2b (5 µg/ml) from Sigma-Aldrich and rabbit anti-ankyrinG (2.5 µg/ml), from Synaptic System (#386003), both diluted in incubation buffer (1% (w/v) BSA, 0.3% (v/v) Triton X-100 in PB). After extensive washing in incubation buffer, the secondary antibodies, Alexa 488 donkey anti-mouse (2 µg/ml; #715-546-151) and Alexa 594 donkey anti rabbit (2 µg/ml; #711-586-152) from Jackson Immunoresearch were incubated for 2 h at room temperature and then washed. After staining, the coverslips were mounted with Vectashield Vibrance (Vector H-1700-10).

### Confocal fluorescence microscopy and image analysis

Images were acquired on a confocal laser scanning microscope (LSM 780 Zeiss) using the same settings to compare intensities between deprived and open sides after 4 days of MD. All images were acquired using a Plan-Apo 63X /1.4 Oil-immersion objective lens. All confocal images were acquired at 0.38-μm z axis steps and with a 1,024 × 1,024-pixel resolution. Photons emitted from the two dyes were detected sequentially with one photomultiplier tube for each dye to minimize cross-talk. Images were prepared using Adobe Photoshop.

Analysis was performed using the ImageJ software (NIH). Images stacks were converted into single maximum intensity z axis projections. AIS was identified by AnkyrinG staining and was selected with threshold determination of fluorescent labeling area (red stain). Analysis of ankyrin G and Kv1.1 staining were performed using Fiji-ImageJ software (NIH), as previously shown (Grubb & Burrone, 2010; Zhang *et al*., 2021). Briefly, a line was drawn along the axon, starting near the cell body. Data were smoothed every 1 µm using the Sigma Plot 11 software. AIS start, maximum and end positions were determined. The origin point was the same for ankyrin G and Kv1.1 labeling. To determine AIS length, AIS start and end points were defined in reference to ankyrin G maximum intensity (Zhang *et al*., 2021). The fluorescence intensity values for Kv1.1 channel were obtained by averaging the intensity over the total length of the measured Kv1.1 signal.

### Statistics

Pooled data are presented as mean ± S.E.M. Statistical analysis was performed using Wilcoxon test, Mann-Whitney *U*-test, or Kruskal-Wallis test. The significant criterion was usually set to p<0.05.

## Acknowledgments

We thank Dr E Carlier for advices in early experiments, Dr B Marquèze-Pouey for help with sutures, Prof. T David for his support and K Milton, O Toutendji & A Venture for excellent animal care.

## Funding

Supported by INSERM, CNRS, AMU, FRM (DVS20131228768 to DomD), ANR (LoGiK, ANR-17-CE16-0022 to DomD, ANR-21-CE16-0013 to DomD & RB), NeuroMarseille (AMX-19-IET-004), and Grant PID2024-159155OB-I00 to JJG, funded by MCIN / AEI / https://doi.org/10.13039/501100011033. This work also received support from the French government under the “France 2030” investment plan, as part of the Initiative d’Excellence d’Aix-Marseille Université – A*MIDEX, AMX-22-RE-AB-187 (to DomD) and AMX-22-RE-V2-007 (to DomD).

## Author contributions

A.A. performed the experiments, analysed the data and wrote the manuscript; L.F.M. performed the experiments and analysed the data and statistics; C.I.B. performed the experiments and analysed the data; MD performed the experiments and analysed the data; EZ performed the experiments; DanD analysed the data; JJG analysed the data and statistics; RB analysed the data, and wrote the manuscript; MR designed the research, initiated and performed experiments, analysed the data and statistics, and edited the manuscript; DomD designed the research, provided funding, analysed the data and statistics, built figures and wrote the manuscript.

## Competing Interests

The authors declare no competing interests.

## Data and Materials Availability

All data needed to evaluate the conclusions in the paper are present in the paper and/or the Supplementary Materials.

---

**Figure S1.**
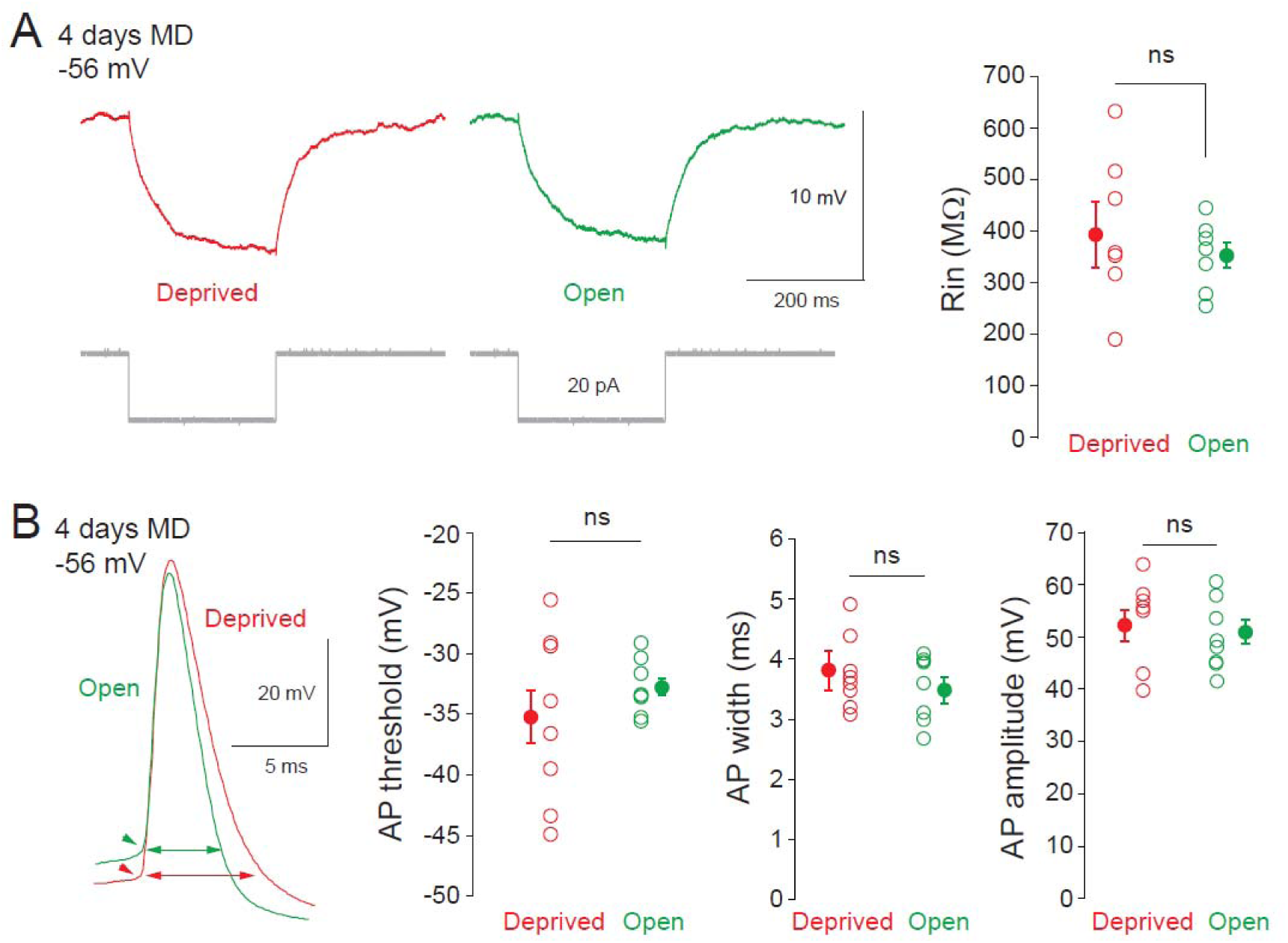
Rin and AP threshold are comparable after 4 days of MD. A. Comparison of input resistance (Rin) and capacitance in neurons recorded in the deprived (red) and open (green) regions of the dLGN. ns, Mann-Whitney U-test, p>0.1. B. Comparison of AP threshold, AP width, AP amplitude in neurons recorded in the deprived (red) and open (green) regions of the dLGN. ns, Mann-Whitney U-test, p>0.1.

**Figure S2.**
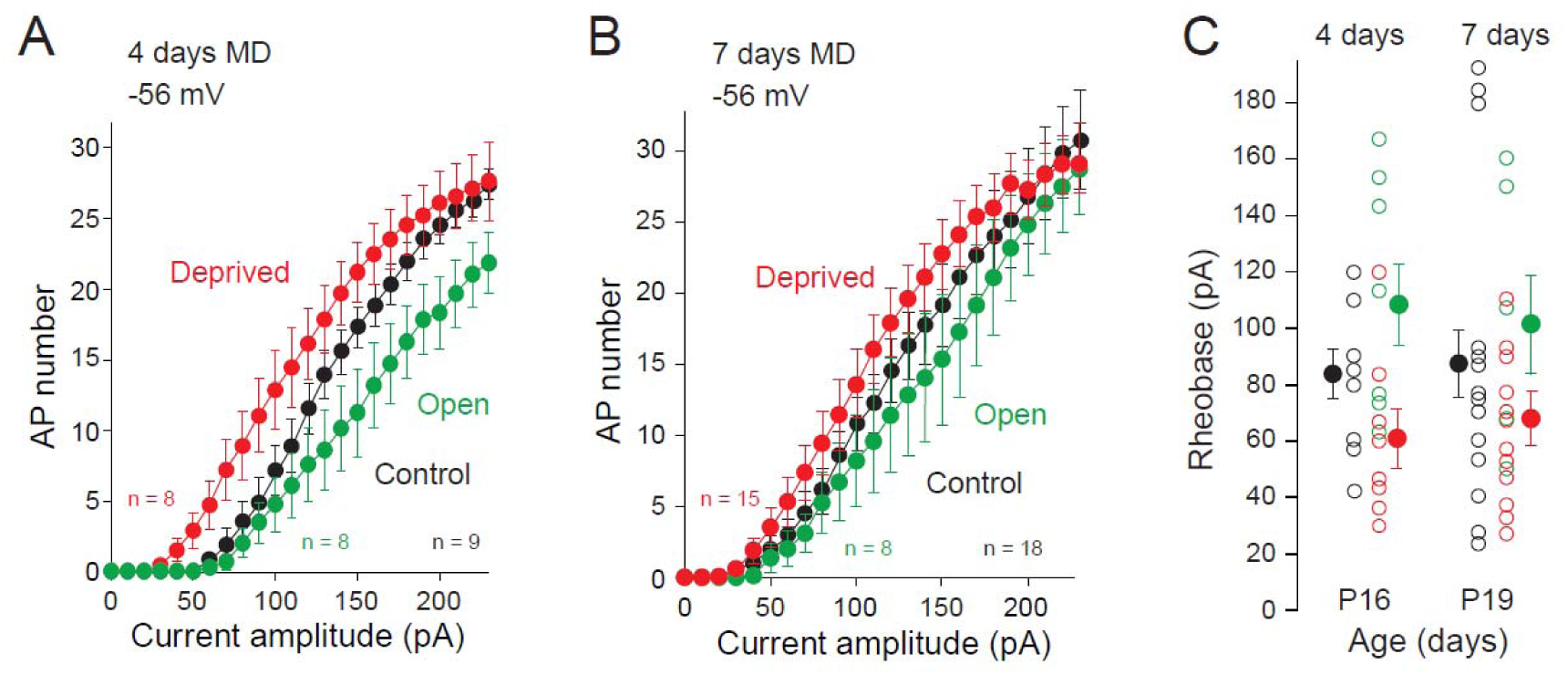
Non deprived animals display an intermediate IE. A. Input-output curves of neurons recorded in control animals at P16 (black dots), deprived (red) or open (green) following 4 days starting at P12 (i.e., in P16 rats). B. Input-output curves of neurons recorded in control animals at P19 (black dots), deprived (red) or open (green) following 7 days starting at P12 (i.e., in P19 rats). C. Comparison of the dLGN neuron rheobase in control animals versus in deprived or open regions. Note that the controls are on average in between the deprived and open means.

**Figure S3.**
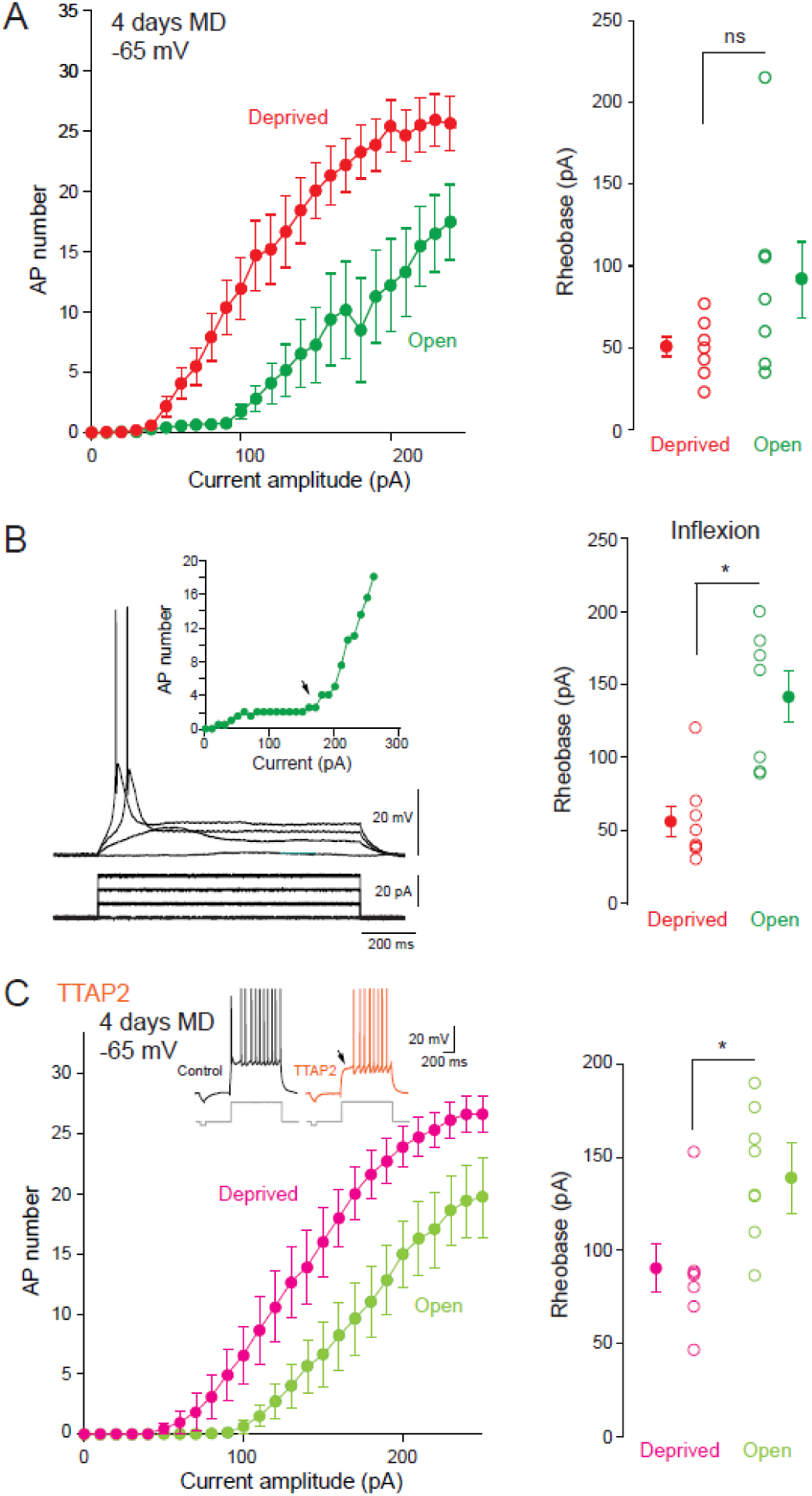
Homeostatic change in IE measured in dLGN neurons at −65 mV after 4 days of MD. A. Left, input-output curves of neurons from the deprived (red) and open (green) zone of the dLGN. Right, comparison of the rheobase in each case. ns, Mann-Whitney U-test, p>0.1. B. Contribution of the T-type calcium potential in the apparent rheobase measured at −65 mV. Left, representative traces showing the T-type calcium potential evoked by the depolarizing current. Right, comparison of the rheobase in deprived and open zone of the dLGN. *, Mann-Whitney U-test, p<0.05. C. In the presence of the T-type calcium blocker, TTA-P2, the initial bust is absent and a significant difference in the rheobase is observed. *, Mann-Whitney U-test, p<0.05.

**Figure S4.**
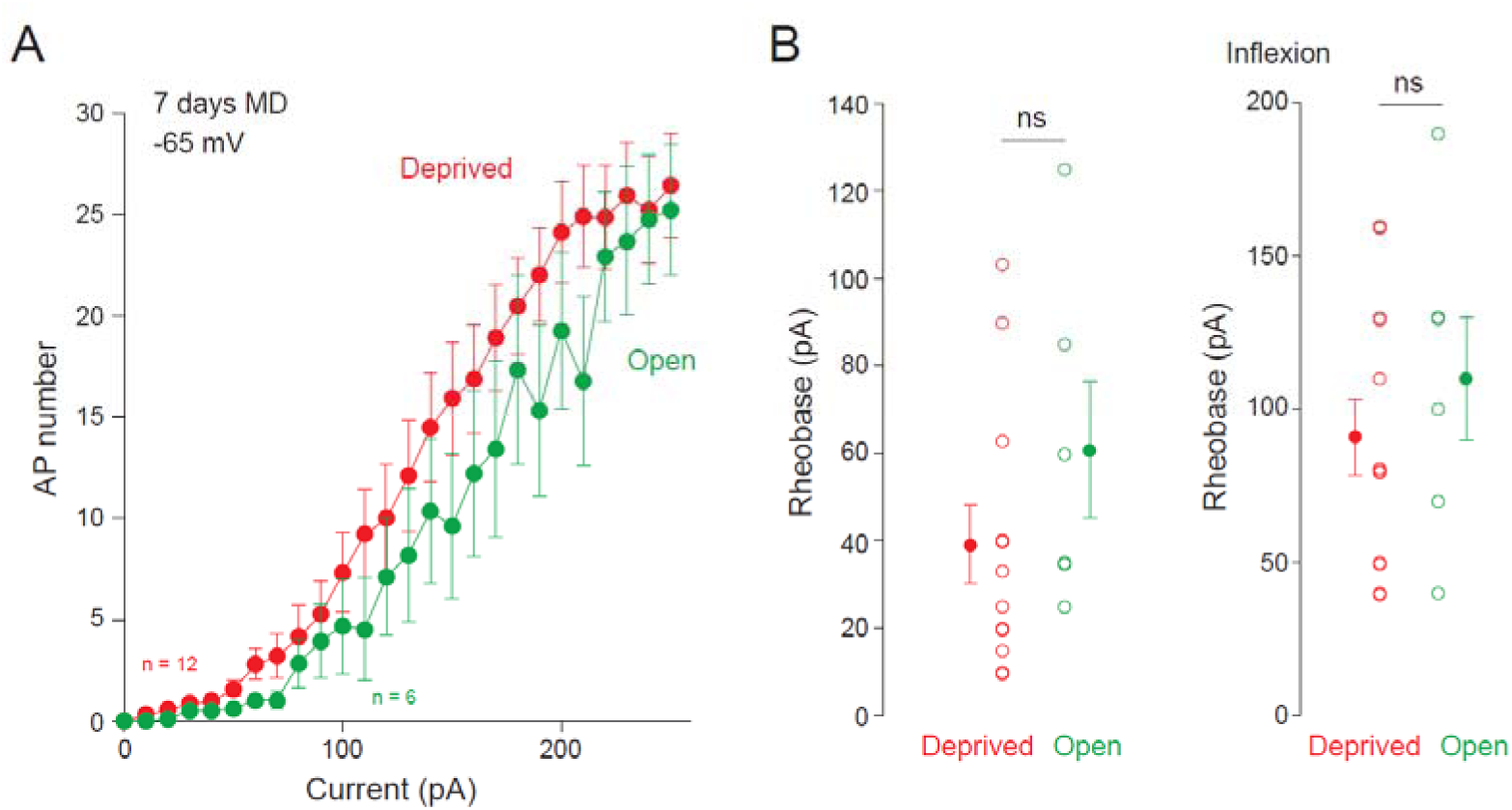
No change in excitability after 7 days of MD at −65 mV. A. Input-output curves of neurons recorded at −65 mV in the deprived (red) or open (green) sides of the dLGN after 7 days of MD. B. Analysis of the rheobase. Left, raw data including the rheobase due to the T-type calcium potential. Right, rheobase measured at the inflexion point. ns, Mann-Whitney U-test, p>0.1.

**Figure S5.**
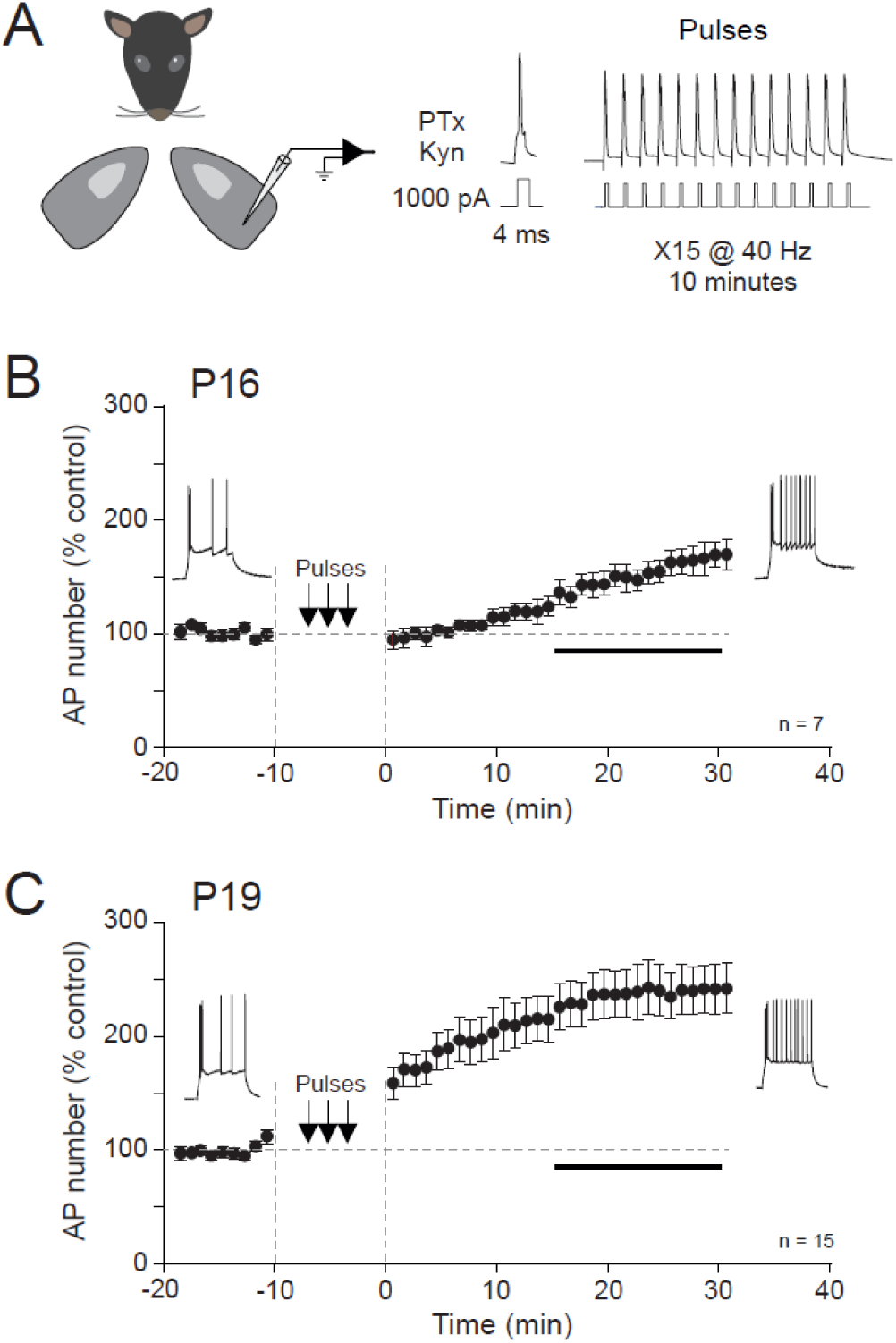
LTP-IE at P16 and P19 in control animals. A. Induction protocol of LTP-IE in dLGN neurons from control animals (both eyes open). B. LTP-IE induced by spiking activity during 10 minutes in P16 animals. C. LTP-IE induced by spiking activity during 10 minutes in P19 animals. Note the larger amplitude of LTP-IE at P19 compared to P16.

**Figure S6.**
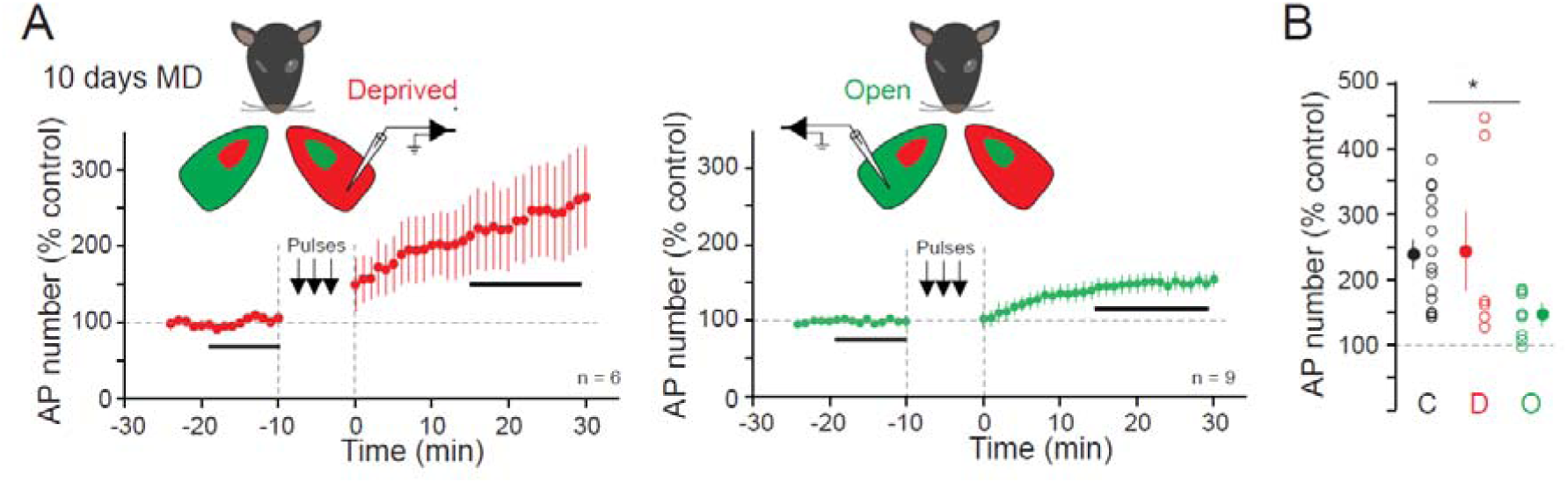
Reduction of LTP-IE in open neurons after 10 days of MD. A. LTP-IE in neurons activated by the deprived eye (left) and open eye (right) after 10 days of MD (data from Duménieu et al., 2025). B. Comparison of the magnitude of LTP-IE in control, deprived and open neurons (data from Duménieu et al., 2025). *, Mann-Whitney U-test, p<0.05.

**Figure S7.**
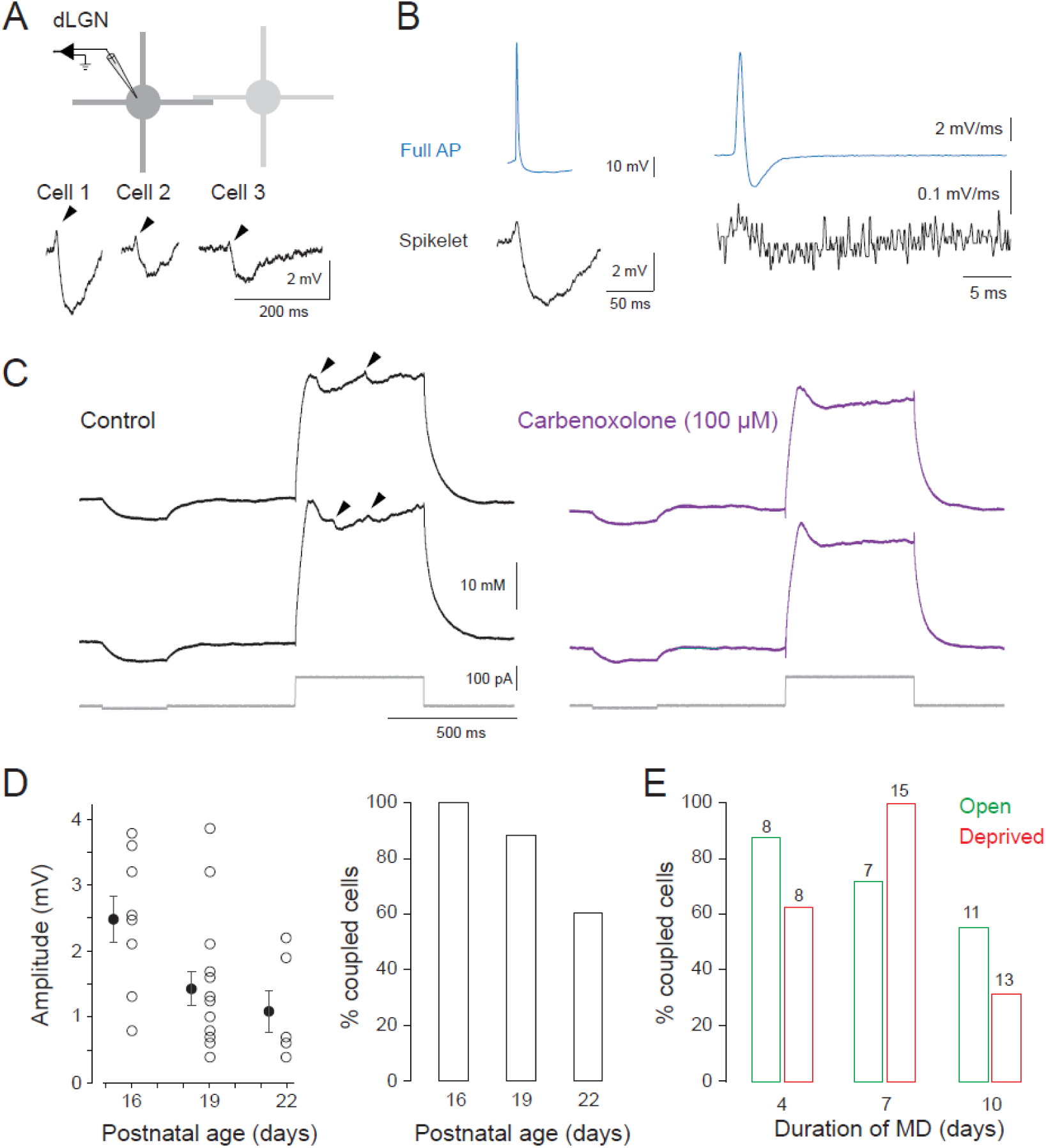
Electrical coupling. A. Spikelet preceding the attenuated AHP in three different dLGN neurons. Arrowheads signal the electrotonically attenuated action potentials. B. Comparison of the dV/dt of a full action potential evoked in the Cell 1 (top trace) with those of the spikelet recorded in Cell 1. C. Carbenoxolone suppresses electrical coupling in dLGN neuron. D. Developmental decline of the amplitude (left) and % of coupled cells (right) in control animals (i.e., with binocular vision). E. Proportion of electrical coupling in deprived (red) and open (green) dLGN neurons following MD. Note the difference in the proportion of coupled cells at 10 days of MD.

**Figure S8.**
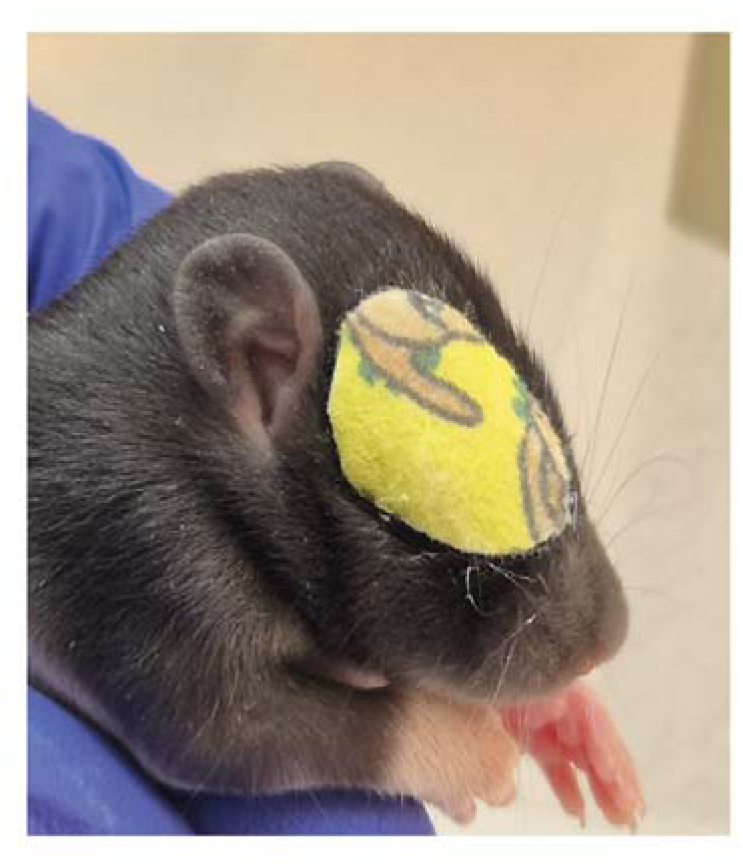
Eye-patch secures the eyelid sutures. Picture of a P12 Long Evans rat wearing a portion of eye-patch glued onto the sutured eyelids to delay the removal of the sutures with the claws.

## Notes

### Competing Interest Statement

The authors have declared no competing interest.

## References

Alcamí P & Pereda AE (2019). Beyond plasticity: the dynamic impact of electrical synapses on neural circuits. Nat Rev Neurosci 20, 253–271.

Bhandari A, Ward TW, Smith J & Van Hook MJ (2022). Structural and Functional Plasticity in the Dorsolateral Geniculate Nucleus of Mice following Bilateral Enucleation. Neuroscience 488, 44–59.

Binda P, Kurzawski JW, Lunghi C, Biagi L, Tosetti M & Morrone MC (2018). Response to short-term deprivation of the human adult visual cortex measured with 7T BOLD ed. Pasternak T & Gold JI. eLife 7, e40014.

Cudmore RH, Fronzaroli-Molinieres L, Giraud P & Debanne D (2010). Spike-time precision and network synchrony are controlled by the homeostatic regulation of the D-type potassium current. J Neurosci 30, 12885–12895.

Desai NS, Rutherford LC & Turrigiano GG (1999). Plasticity in the intrinsic excitability of cortical pyramidal neurons. Nat Neurosci 2, 515–520.

Dugué GP, Brunel N, Hakim V, Schwartz E, Chat M, Lévesque M, Courtemanche R, Léna C & Dieudonné S (2009). Electrical coupling mediates tunable low-frequency oscillations and resonance in the cerebellar Golgi cell network. Neuron 61, 126–139.

Duménieu M, Fronzaroli-Molinieres L, Naudin L, Iborra-Bonnaure C, Wakade A, Zanin E, Aziz A, Ankri N, Incontro S, Denis D, Marquèze-Pouey B, Brette R, Debanne D & Russier M (2025). Visual activity enhances neuronal excitability in thalamic relay neurons. Sci Adv 11, eadp4627.

Frégnac Y, Shulz D, Thorpe S & Bienenstock E (1988). A cellular analogue of visual cortical plasticity. Nature 333, 367–370.

Fricker B, Heckman E, Cunningham PC, Wang H & Haas JS (2021). Activity-dependent long-term potentiation of electrical synapses in the mammalian thalamus. J Neurophysiol 125, 476–488.

Galarreta M & Hestrin S (2001). Spike transmission and synchrony detection in networks of GABAergic interneurons. Science 292, 2295–2299.

Garrido JJ, Giraud P, Carlier E, Fernandes F, Moussif A, Fache M-P, Debanne D & Dargent B (2003). A targeting motif involved in sodium channel clustering at the axonal initial segment. Science 300, 2091–2094.

Gasselin C, Inglebert Y & Debanne D (2015). Homeostatic regulation of h-conductance controls intrinsic excitability and stabilizes the threshold for synaptic modification in CA1 neurons. J Physiol (Lond) 593, 4855–4869.

Grubb MS & Burrone J (2010). Activity-dependent relocation of the axon initial segment fine-tunes neuronal excitability. Nature 465, 1070–1074.

Haas JS, Zavala B & Landisman CE (2011). Activity-dependent long-term depression of electrical synapses. Science 334, 389–393.

Holmes JM & Clarke MP (2006). Amblyopia. Lancet 367, 1343–1351.

Hubel DH & Wiesel TN (1970). The period of susceptibility to the physiological effects of unilateral eye closure in kittens. J Physiol 206, 419–436.

Hughes S, Lőrincz M, Turmaine M & Crunelli V (2011). Thalamic Gap Junctions Control Local Neuronal Synchrony and Influence Macroscopic Oscillation Amplitude during EEG Alpha Rhythms. Front Psychol; DOI: 10.3389/fpsyg.2011.00193.

Hughes SW, Lörincz M, Cope DW, Blethyn KL, Kékesi KA, Parri HR, Juhász G & Crunelli V (2004). Synchronized oscillations at alpha and theta frequencies in the lateral geniculate nucleus. Neuron 42, 253–268.

Ibata K, Sun Q & Turrigiano GG (2008). Rapid synaptic scaling induced by changes in postsynaptic firing. Neuron 57, 819–826.

Jaepel J, Hübener M, Bonhoeffer T & Rose T (2017). Lateral geniculate neurons projecting to primary visual cortex show ocular dominance plasticity in adult mice. Nat Neurosci 20, 1708–1714.

Jamann N, Dannehl D, Lehmann N, Wagener R, Thielemann C, Schultz C, Staiger J, Kole MHP & Engelhardt M (2021). Sensory input drives rapid homeostatic scaling of the axon initial segment in mouse barrel cortex. Nat Commun 12, 23.

Jenkins PM & Bender KJ (2025). Axon initial segment structure and function in health and disease. Physiol Rev 105, 765–801.

Karmarkar UR & Buonomano DV (2006). Different forms of homeostatic plasticity are engaged with distinct temporal profiles. Eur J Neurosci 23, 1575–1584.

Kepler TB, Marder E & Abbott LF (1990). The Effect of Electrical Coupling on the Frequency of Model Neuronal Oscillators. Science 248, 83–85.

Kleinschmidt A, Bear MF & Singer W (1987). Blockade of “NMDA” receptors disrupts experience-dependent plasticity of kitten striate cortex. Science 238, 355–358.

Krahe TE & Guido W (2011). Homeostatic plasticity in the visual thalamus by monocular deprivation. J Neurosci 31, 6842–6849.

Krosigk von M, Bal T & McCormick DA (1993). Cellular Mechanisms of a Synchronized Oscillation in the Thalamus. Science 261, 361–364.

Kuba H, Yamada R, Ishiguro G & Adachi R (2015). Redistribution of Kv1 and Kv7 enhances neuronal excitability during structural axon initial segment plasticity. Nat Commun 6, 8815.

Landisman CE & Connors BW (2005). Long-term modulation of electrical synapses in the mammalian thalamus. Science 310, 1809–1813.

Lee S-C, Cruikshank SJ & Connors BW (2010). Electrical and chemical synapses between relay neurons in developing thalamus. J Physiol 588, 2403–2415.

Lunghi C, Berchicci M, Morrone MC & Di Russo F (2015). Short-term monocular deprivation alters early components of visual evoked potentials. J Physiol 593, 4361–4372.

Lunghi C, Burr DC & Morrone C (2011). Brief periods of monocular deprivation disrupt ocular balance in human adult visual cortex. Curr Biol 21, R538–539.

Maffei A, Nelson SB & Turrigiano GG (2004). Selective reconfiguration of layer 4 visual cortical circuitry by visual deprivation. Nat Neurosci 7, 1353–1359.

Marder E & Goaillard J-M (2006). Variability, compensation and homeostasis in neuron and network function. Nat Rev Neurosci 7, 563–574.

Morgan PJ, Bourboulou R, Filippi C, Koenig-Gambini J & Epsztein J (2019). Kv1.1 contributes to a rapid homeostatic plasticity of intrinsic excitability in CA1 pyramidal neurons in vivo. Elife 8, e49915.

Morishita H & Hensch TK (2008). Critical period revisited: impact on vision. Curr Opin Neurobiol 18, 101–107.

Mrsic-Flogel TD, Hofer SB, Ohki K, Reid RC, Bonhoeffer T & Hübener M (2007). Homeostatic regulation of eye-specific responses in visual cortex during ocular dominance plasticity. Neuron 54, 961–972.

Nataraj K, Le Roux N, Nahmani M, Lefort S & Turrigiano G (2010). Visual deprivation suppresses L5 pyramidal neuron excitability by preventing the induction of intrinsic plasticity. Neuron 68, 750–762.

Riegle KC & Meyer RL (2007). Rapid homeostatic plasticity in the intact adult visual system. J Neurosci 27, 10556–10567.

Rose T & Bonhoeffer T (2018). Experience-dependent plasticity in the lateral geniculate nucleus. Curr Opin Neurobiol 53, 22–28.

Russier M, Duménieu M, Wakade A, Incontro S, Fronzaroli-Molinieres L & Debanne D (2024). Inducing Long-Term Plasticity of Intrinsic Neuronal Excitability in Neurons of the Dorsal Lateral Geniculate Nucleus. J Vis Exp; DOI: 10.3791/65950.

Sevetson J, Fittro S, Heckman E & Haas JS (2017). A calcium-dependent pathway underlies activity-dependent plasticity of electrical synapses in the thalamic reticular nucleus. J Physiol 595, 4417–4430.

Sommeijer J-P, Ahmadlou M, Saiepour MH, Seignette K, Min R, Heimel JA & Levelt CN (2017). Thalamic inhibition regulates critical-period plasticity in visual cortex and thalamus. Nat Neurosci 20, 1715–1721.

Steriade M, McCormick DA & Sejnowski TJ (1993). Thalamocortical oscillations in the sleeping and aroused brain. Science 262, 679–685.

Storchi R, Bedford RA, Martial FP, Allen AE, Wynne J, Montemurro MA, Petersen RS & Lucas RJ (2017). Modulation of Fast Narrowband Oscillations in the Mouse Retina and dLGN According to Background Light Intensity. Neuron 93, 299–307.

Turrigiano GG (2017). The dialectic of Hebb and homeostasis. Philos Trans R Soc Lond B Biol Sci 372, 20160258.

Turrigiano GG & Nelson SB (2004). Homeostatic plasticity in the developing nervous system. Nat Rev Neurosci 5, 97–107.

Van Hook MJ, Monaco C, Bierlein ER & Smith JC (2020). Neuronal and Synaptic Plasticity in the Visual Thalamus in Mouse Models of Glaucoma. Front Cell Neurosci 14, 626056.

Vaughn MJ, Yellamelli N, Burger RM & Haas JS (2025). Dopamine receptors D1, D2, and D4 modulate electrical synapses and excitability in the thalamic reticular nucleus. J Neurophysiol 133, 374–387.

Weyand TG, Boudreaux M & Guido W (2001). Burst and tonic response modes in thalamic neurons during sleep and wakefulness. J Neurophysiol 85, 1107–1118.

Wiesel TN & Hubel DH (1963). SINGLE-CELL RESPONSES IN STRIATE CORTEX OF KITTENS DEPRIVED OF VISION IN ONE EYE. J Neurophysiol 26, 1003–1017.

Zbili M, Rama S, Benitez M-J, Fronzaroli-Molinieres L, Bialowas A, Boumedine-Guignon N, Garrido JJ & Debanne D (2021). Homeostatic regulation of axonal Kv1.1 channels accounts for both synaptic and intrinsic modifications in the hippocampal CA3 circuit. Proc Natl Acad Sci U S A 118, e2110601118.

Zhang W, Ciorraga M, Mendez P, Retana D, Boumedine-Guignon N, Achón B, Russier M, Debanne D & Garrido JJ (2021). Formin Activity and mDia1 Contribute to Maintain Axon Initial Segment Composition and Structure. Mol Neurobiol 58, 6153–6169.

Zsiros V & Maccaferri G (2005). Electrical coupling between interneurons with different excitable properties in the stratum lacunosum-moleculare of the juvenile CA1 rat hippocampus. J Neurosci 25, 8686–8695.

